## Supplemental table 1 and table 2 for "Physiological oxygen concentration during sympathetic primary neuron culture improves neuronal health and reduces HSV-1 reactivation"

**Supplemental tables**

**Table S1: Cell Body Scoring Index**

| Score | Description |
| --- | --- |
| 0 | Large, phase bright cell bodies. Clear with no fragmentation or vesiculation. |
| 1 | Small, phase bright cell bodies. Clear with no fragmentation or vesiculation. |
| 2 | Cell bodies do not have fragmentation but are not phase bright. Sometimes appear transparent.^1^ |
| 3 | Cell bodies with fragmentation but few dead neurons or corpses |
| 4 | Cell bodies with fragmentation with many corpses present and neurons starting to detach |
| 5 | Complete cell death. Neurons detached. |

**Table S2: Axon Scoring Index**

| Score | Description |
| --- | --- |
| 0 | Axons totally smooth with no blebbing or fragmentation. Branched and form a spider web-like network |
| 1 | Axons smooth but grow straight |
| 2 | Blebbing on the axons but no apparent fragmentation |
| 3 | Fragmentation starting to appear in <50% of the neurons |
| 4 | Fragmentation in >50% of the neurons |
| 5 | No axons remaining |
